# Conserved role for spliceosomal component PRPF40A in microexon splicing

**DOI:** 10.1101/2024.09.26.615222

**Authors:** Bikash Choudhary, Adam Norris

## Abstract

Microexons (exons ≤30 nts) are important features of neuronal transcriptomes, but pose mechanistic challenges to the splicing machinery. We previously showed that PRP-40, a component of the U1 spliceosome, is globally required for microexon splicing in *C. elegans*. Here we show that the homologous PRPF40A is also globally required for microexon splicing in mouse neuroblastoma cells. We find that PRPF40A co-regulates microexons along with SRRM4, a neuron-specific regulator of microexon splicing. The relationship between exon size and dependence on PRPF40A/SRRM4 is distinct, with SRRM4-dependence exhibiting a size threshold (∼30 nts) and PRPF40A-dependence exhibiting a graded decrease as exon size increases. Finally, we show that PRPF40A knockdown causes an increase in productive splicing of its spliceosomal binding partner *Luc7l* by skipping of a small “poison exon.” Similar homeostatic cross-regulation is often observed across paralogous RNA binding proteins. Here we find this concept likewise applies across evolutionarily unrelated but functionally and physically coupled spliceosomal components.

## INTRODUCTION

Many eukaryotic genes are split into protein-coding exons interrupted by intervening introns. These introns are removed by the spliceosome to generate mature mRNAs. While intron sizes vary substantially across genes and species, exon size is more constrained, with a median size of ∼120 nucleotides (nts) across various animal species^1^. Indeed, exons below ∼51 nts are spliced inefficiently, partly due to physical constraints impeding spliceosomal assembly flanking small exons ^2,3^

Nevertheless, a subset of unusually small exons referred to as microexons, often defined as exons ≤30 nts, has recently emerged as a distinct functional class. Microexons are enriched in neuronal genes, and tend to be alternatively spliced such that the microexon is included in neurons but skipped in other cells^4–6^. Moreover, dysregulated microexon splicing is as a common molecular feature in autism spectrum disorder^5^. Thus, in spite of mechanistic hurdles complicating the splicing of exons below a certain size, microexons constitute an important feature the neuronal transcriptome.

Studies on the regulation of microexon splicing have identified a handful of required factors. The splicing factors RBFOX and PTBP1 were shown to regulate a number of microexons in mouse cells ^4^, and the neuronal splicing factor SRRM4 (aka nSR100) was identified as a factor required for global micrexon splicing in mice and human cells^5,7,8^. Likewise, we recently found in *C. elegans* that PRP-40, a component of the U1 snRNP, is globally required for microexon splicing^9^.

The potential mechanistic relationships among these observations remains to be determined. In particular, we wanted to investigate the nature of global microexon regulators SRRM4 and PRP-40: is PRP-40 required globally for microexon splicing in mouse as it is in *C. elegans*? If so, are the attributes of PRP-40-dependent exons and SRRM4-dependent exons identical? The possibility of different microexon regulatory strategies between mouse and worm is raised by evidence that mouse SRRM4 stimulates microexon splicing via its C-terminal domain, while *C. elegans* does not encode such a domain in its closest protein homologue (RSR-2)^10^. We therefore set out to determine in mouse cells the relationship between PRPF40A and SRRM4 in microexon splicing.

We find that knockdown of either PRPF40A or SRRM4 in mouse neuroblastoma cells causes a global decrease in microexon inclusion, thus the activity of PRPF40A in mouse is similar to PRP-40 in worm. Morerover, mouse microexons require both PRPF40A and SSRM4 simultaneously for efficient splicing. However, PRPF40A-dependent exons demonstrate a graded decrease in PRPF40A dependence as size increases, while SRRM4-dependent exons display size threshold, with exons above a ∼30 nucleotide threshold not requiring SRRM4. Additionally, we reveal that PRPF40A knockdown causes a large increase of productive splicing of its spliceosomal binding partner *Luc7l* by skipping a small poison exon. We propose that this is similar to cases of homeostatic compensatory regulation across paralagous RNA binding proteins, thus extending the concept to compensatory regulation across two evolutionarily unrelated but structurally neighboring factors (PRPF40A and LUC7L).

## RESULTS

### PRPF40A Knockdown Causes Global Loss of Microexon Splicing

PRP40 is a component of the U1 snRNP^11^. We previously showed that loss of *C. elegans prp-40* does not result in systemic splicing defects, as might be expected for a spliceosomal component, but rather results specifically in loss of microexon and small-exon splicing^9^. Yeast and worm genomes encode a single PRP40 gene, while mouse and human genomes encode two paralagous genes, PRPF40A and PRPF40B, with PRPF40A most resembling the PRP40 genes of yeast and worms (Fig 1A). We showed that, as in worms, knockdown of PRPF40A in mouse neuroblastoma cells results in loss of microexon splicing for the handful of exons we tested^9^. Here we wanted to extend these findings to a global analysis of splicing controlled by PRPF40A. To this end we performed PRPF40A knockdown in mouse neuroblastoma N2a cells followed by RNA-Seq.

**Figure 1.**
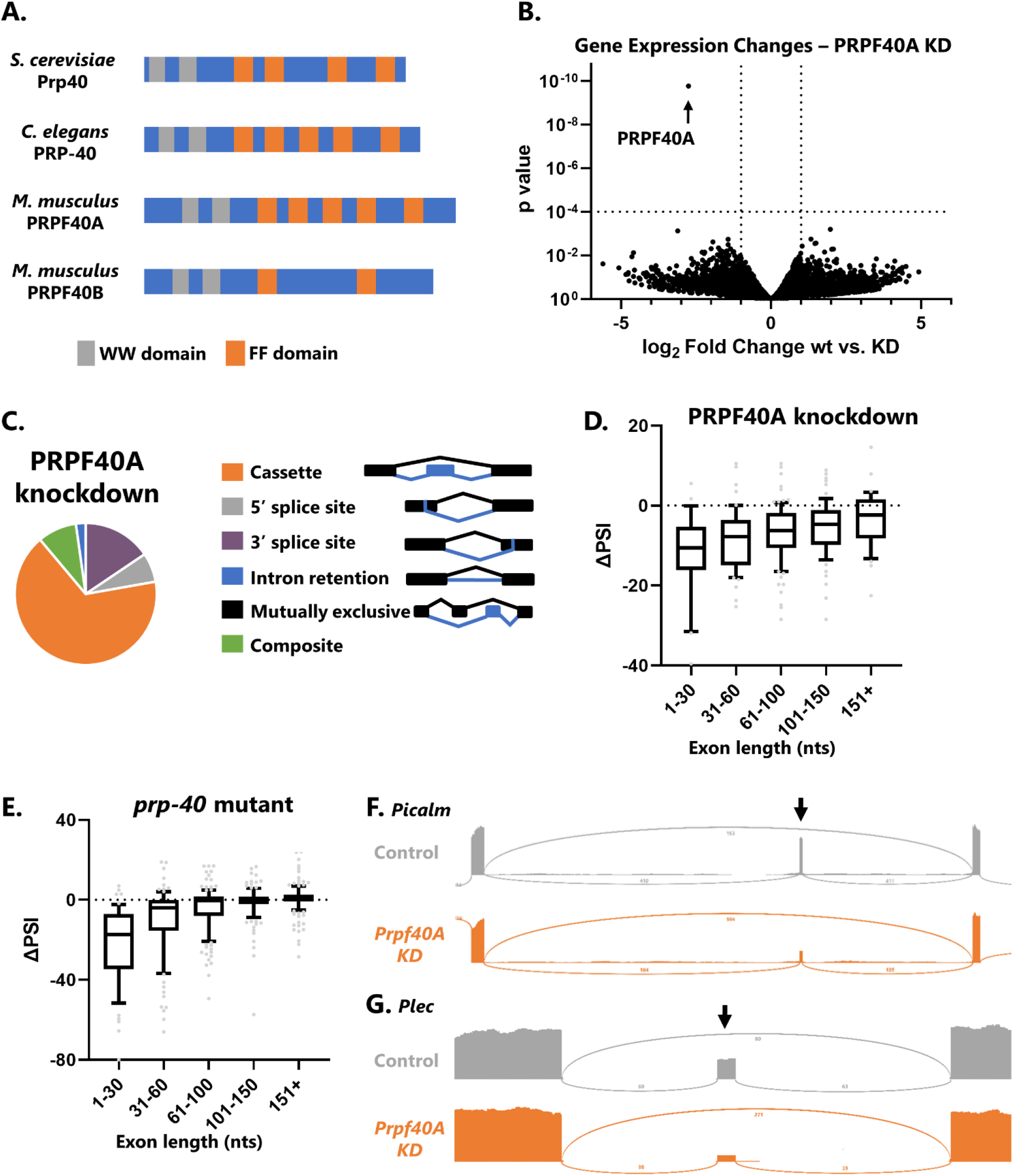
PRPF40A knockdown causes global loss of microexon and small exon splicing. (A) Protein domain structure of PRP-40 homologues, modeled after TreeFam database^12^. (B) Volcano plot showing changes in gene expression as determined by DESeq2 upon PRPF40A siRNA knockdown. (C) The effect of PRPF40A knockdown on alternative splicing is largely restricted to changes in cassette exon inclusion (67%). (D) Effects of PRPF40A knockdown on cassette exon inclusion binned by exon size, including microexons (1-30 nts). ΔPSI = change in percent spliced in from control to PRPF40A conditions. (E) Same as in D, except for *C. elegans prp-40* loss-of-function mutants. (F-G) Sashimi plots showing that PRPF40A knockdown results in dramatic decreases in microexon inclusion, but does not affect splicing of upstream or downstream exons, or lead to increased retention of the flanking introns.

Knockdown of PRPF40A causes little change in global gene expression levels except for, as expected, a strong decrease in PRPF40A levels (Fig 1B). PRPF40A knockdown has specific effects on alternative splicing, largely limited to dysregulated cassette exon splicing (aka exon skipping). Other types of alternative splicing, for example intron retention or 3’ splice site selection, are largely unaffected (Fig 1C). Moreover, the effect of PRPF40A is directional, as knockdown consistently results in negative changes in Percent Spliced In (ΔPSI, Fig 1D), indicating that the role of PRPF40A is to facilitate cassette exon inclusion.

The effect of PRPF40A on cassette exon inclusion depends on exon length. Microexons (≤30 nts) are most strongly affected, and as exon size increases, the dependence on PRPF40A decreases (Fig 1D). This pattern of size dependence is similar to our previous observations in *C. elegans prp-40* mutants (Fig 1E). Therefore, mouse PRPF40A, like *C. elegans* PRP-40, is required specifically for the inclusion of microexons and small exons. Furthermore, PRPF40A knockdown does not lead to an increase in retention of the introns flanking microexons (examples in Fig 1F-G). This indicates that PRPF40A is not required generally for splicing fidelity, but is required specifically for the splicing decision of microexon inclusion versus skipping.

### SRRM4 and PRPF40A are both globally required for microexons, but with different regulatory features

In parallel with our PRPF40A knockdown we also knocked down two related factors of interest: PRPF40B, the paralog of PRPF40A; and SRRM4, a neuronal splicing factor required for inclusion of many microexons^7^. The results of PRPF40B knockdown are in striking contrast to those of PRPF40A, showing no global relationship between exon size and PRPF40B requirement (Fig 2A). This indicates that PRPF40A, but not PRPF40B, acts in a similar manner to *C. elegans* PRP-40 in stimulating microexon inclusion.

**Figure 2.**
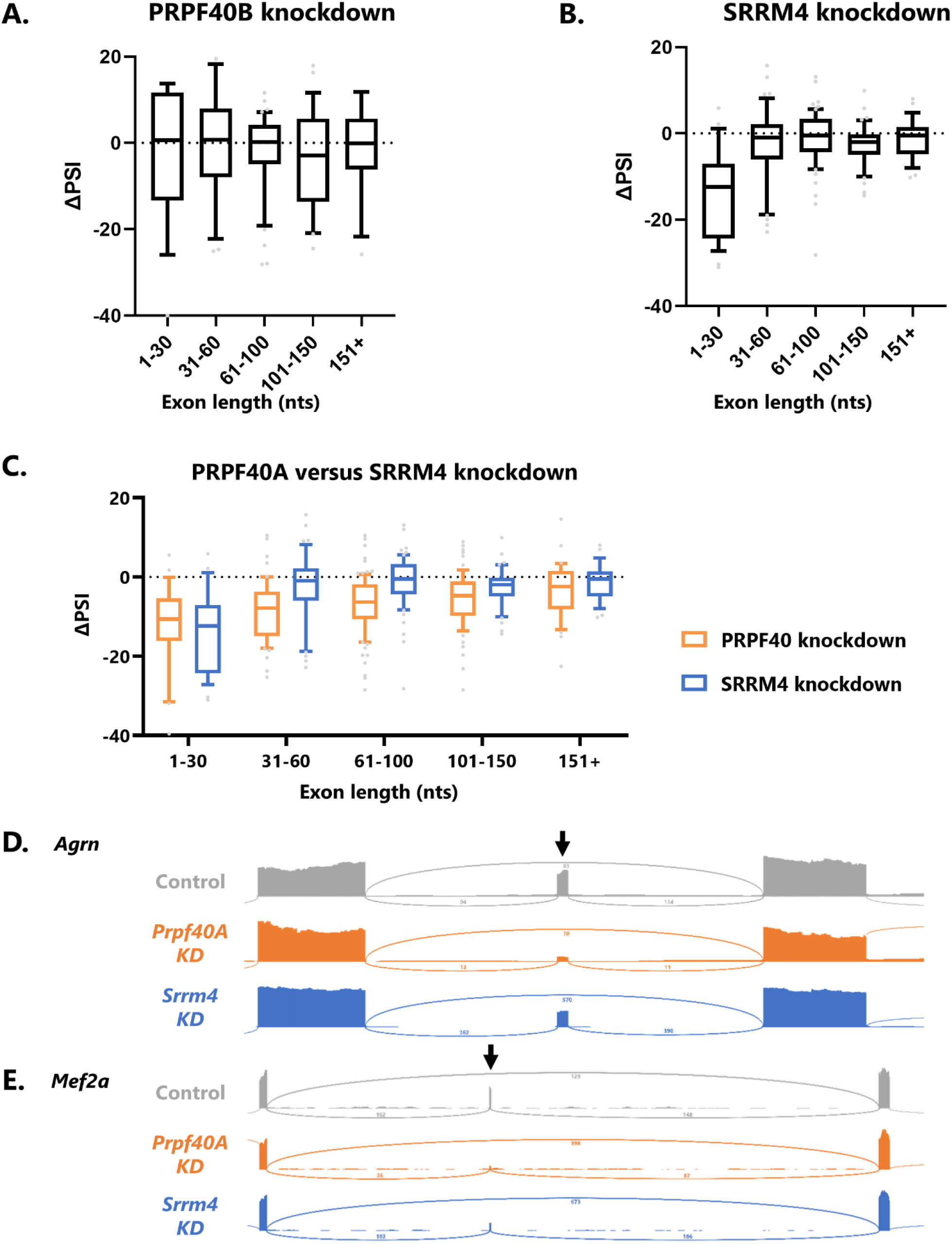
SRRM4 and PRPF40A co-regulate microexons. (A) In contrast with PRPF40A knockdown, PRPF40B knockdown does not result in global splicing defects with respect to exon length. (B) SRRM4 knockdown results in loss of microexon inclusion. (C) Both PRPF40A and SRRM4 are required for microexon splicing, but only PRPF40A is required for other small exon splicing (*e.g*. the 31-60 nt bin). (D) Example of a microexon (in *Agrn*) strongly dependent on PRPF40A but only mildly regulated by SRRM4. (E) Example of a microexon (in *Mef2a)* co-regulated by SRRM4 and PRPF40A.

In contrast, knockdown of SRRM4 results in widespread loss of microexon inclusion, as expected (Fig 2B). The magnitude of microexon dysregulation is similar between SRRM4 and PRPF40A knockdown (Fig 2C). However, the global relationship between exon length and ΔPSI is different for the two factors. PRPF40A-dependent exons show a graded decrease in PRPF40A dependence as their size increases (Fig 2C). But SRRM4-dependent exons display a size threshold, with microexons ≤30 nts strongly dependent on SRRM4, and exons above this threshold largely unaffected. As such, the global effects of PRPF40A and SRRM4 are similar for microexons, but diverge for small exons above the 30 nt threshold (Fig 2C).

Most PRPF40A-dependent microexons are also SRRM4-dependent, and vice versa, although we did detect a few notable exceptions. For example, a 12-nt microexon in the *Agrn* gene is strongly dependent on PRPF40A, but not on SRRM4 (Fig 2D). However, the most common scenario, as in the 24-nt microexon in *Mef2a* (Fig 2E), is that both factors are required for microexon inclusion, indicating that most microexons are co-regulated by PRPF40A and SRRM4.

### PRPF40A knockdown increases productive alternative splicing of its spliceosomal binding partner LUC7L

We noted a particularly strong effect of PRPF40A, but not SRRM4 or PRPF40B, on the splicing of a small 71 nt alternatively-spliced exon in the *Luc7l* gene. *Luc7l* encodes a U1 spliceosomal component that in yeast physically interacts with Prp40^13^ (Fig 3A). Under normal conditions, the exon is included in more than half of *Luc7l* transcripts (53.3%). Inclusion of this exon causes a frameshift and is predicted to encode a truncated LUC7L protein lacking both conserved zinc finger motifs (Fig 3B), and the mRNA is likely to be an NMD substrate^14^. Therefore, the most common splicing outcome in normal conditions leads to non-functional LUC7L due to inclusion of a “poison exon.” A second common splicing outcome is retention of the downstream and/or upstream introns (12% intron retention). The result of this splicing choice is also predicted to be NMD sensitive and encode a truncated, non-functional protein (Fig 3B). Only the exon-skipped product, which constitutes a mere 35% of the spliced output in control N2a cells, is predicted to be functional.

**Figure 3.**
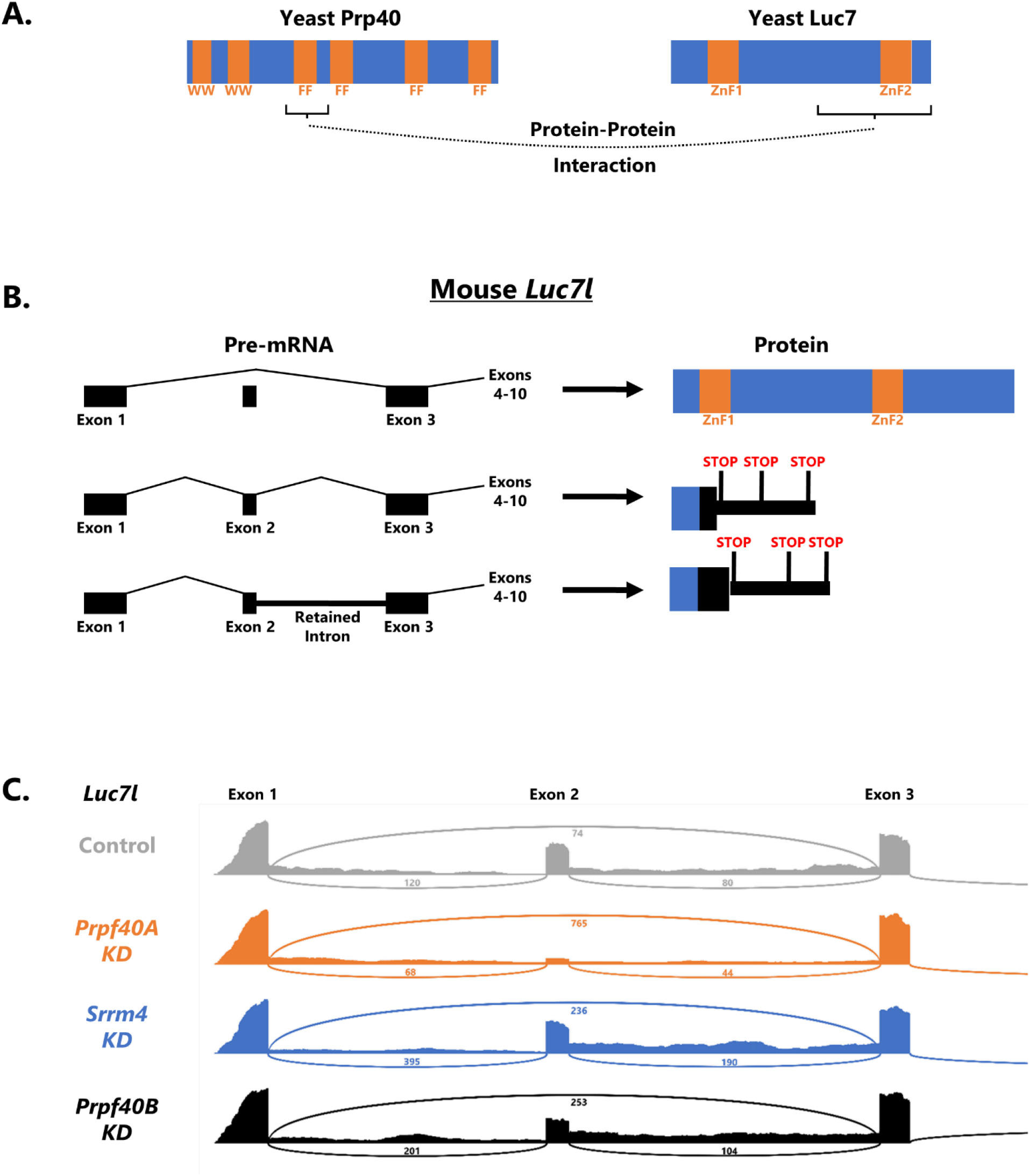
PRPF40A knockdown increases productive splicing of its spliceosomal binding partner LUC7L. (A) Summary of protein-protein interaction data^13^ showing that yeast Prp40 directly interacts with yeast Luc7 via its 1^st^ FF domain. (B) Mouse *Luc7l* gene model showing that if exon 2 is included, or if introns 1 or 2 are retained, then unproductive LUC7L is produced, lacking the region that interacts with Prp40, and containing premature stop codons likely to subject the transcript to NMD. (C) In control N2a cells, the majority of *Luc7l* is spliced into unproductive isoforms. (This is likely an underestimate, given that some unproductive mRNAs are likely destroyed by MMD.) Knockdown of PRPF40A, but not PRPF40B or SRRM4, results in strong increase of the productive isoform (exon 2 skipped and introns 1 and 2 spliced out).

The consequence of PRPF40A knockdown is a dramatic shift in *Luc7l* splicing from unproductive isoforms (exon included and/or intron retained) to the productive exon-skipped isoform: from 35% productive isoform in control to 80% in PRPF40A knockdown (Fig 3C). In the yeast U1 spliceosome, Prp40 and Luc7 are direct protein-protein interactors^13^ and are juxtaposed next to each other in cryoEM structures^11^. As such, a major molecular response to loss of PRPF40A is an increase in its spliceosomal partner LUC7L mediated by productive alternative splicing. A similar response has been observed in human cells upon knockdown of the *Luc7l* paralog *Luc7l2*^14,15^. Knockdown of *Luc7l2* causes an increase in *Luc7l* exon skipping and a decrease in both intron retention and exon inclusion. This response appears to be a compensatory mechanism of the type often seen among paralogous RNA binding proteins^16–18^. Here we extend this phenomenon of compensatory splicing changes to include evolutionarily unrelated but physically associated components of the U1 snRNP. We speculate this might be a mechanism for homeostatic control of spliceosome formation and that abundant LUC7L might partially compensate for loss of its binding partner PRPF40A.

## DISCUSSION

Here we show that the spliceosomal component PRPF40A is globally required for the splicing of microexons and small exons, but not for other types of alternative or constitutive splicing. This role for PRPF40A is shared between mouse and *C. elegans* and thus the specific activity of this spliceosomal component in facilitating microexon inclusion appears to be a deeply conserved phenomenon.

We find that PRPF40A and SRRM4 co-regulate microexon splicing, while small exons (∼31-60 nts) require PRPF40A but not SRRM4. We speculate that for microexons, SRRM4 might directly recruit PRPF40A to facilitate exon inclusion. Affinity-Purification Mass Spectrometry experiments in mouse N2a cells revealed protein-protein interactions between SRRM4 and PRPF40A.^19^ We propose a model (Fig 4) in which SRRM4 binds to microexon-flanking introns via UGC-containing motifs^19^ and recruits PRPF40A to ensure microexon splicing. We previously implicated PRP-40 in an “intron definition” splicing mechanism for microexons whose small size physically precludes them from the typical “exon definition” splicing mechanism favored in many metazoa^2,9^. Thus our model proposes that without SRRM4 and/or PRPF40A, microexons are skipped due to physical inability of the spliceosome to assemble across the exon in an exon definition mechanism. However, in neurons, SRRM4 binds to the flanking intron(s) and recruits PRPF40A, which then facilitates intron-definition-mediated splicing of the flanking introns, thus ensuring microexon inclusion (Fig 4).

**Figure 4.**
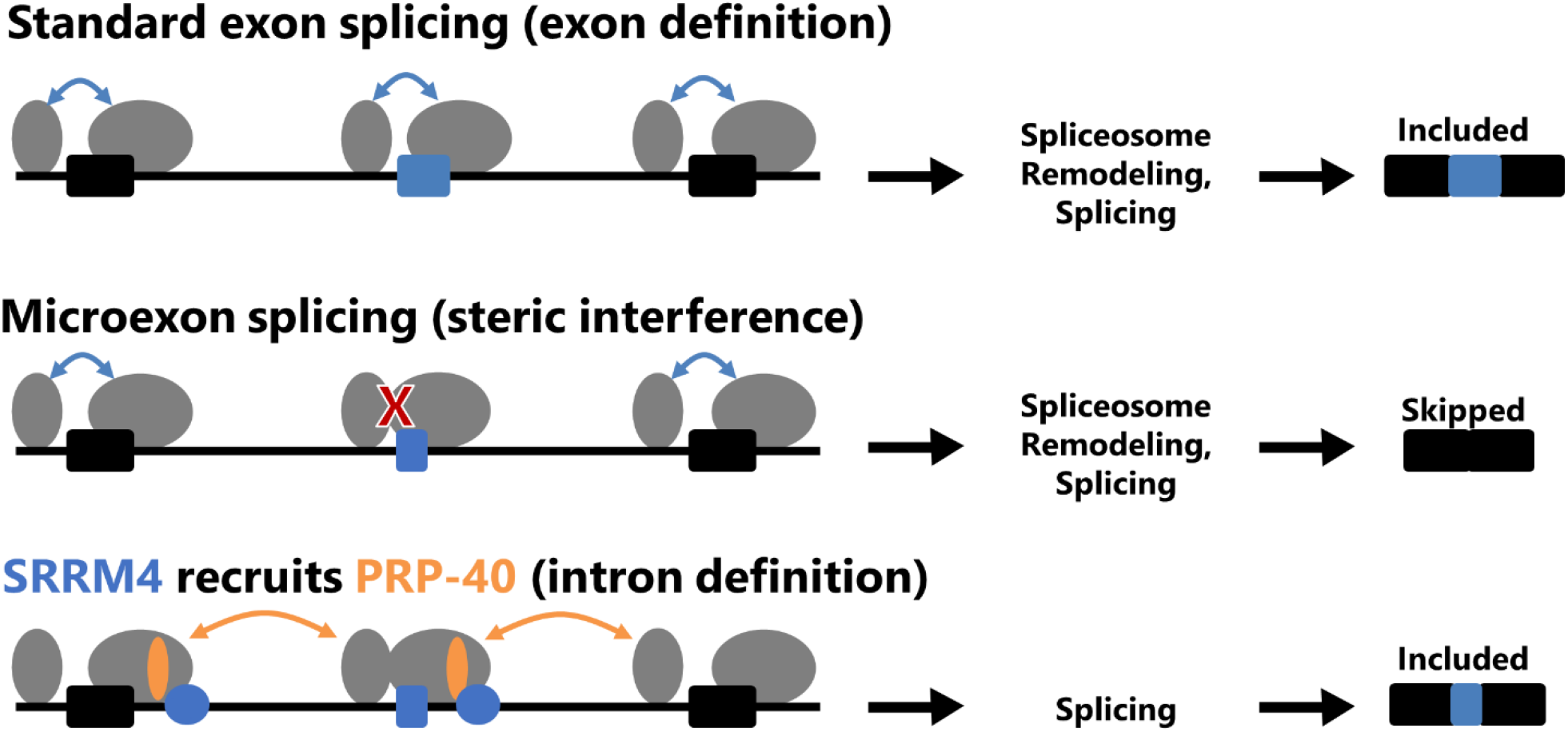
Model for co-regulation of microexons by SRRM4 and PRPF40A. For conventional-sized exons, splicing proceeds via exon definition, in which the unit of recognition is the exon. This is not physically feasible for microexons, which cannot accommodate the binding of both the U1 and U2 snRNPs on such a small sequence space. However, this physical constraint can be overcome when SRRM4, binding to UGC-containing motifs in the flanking intron, recruits PRPF40A, which then facilitates intron definition-mediated splicing of the flanking introns, leading to microexon inclusion in the mature mRNA.

While the activities of PRPF40A and PRP-40 are similar between mouse and worm, there is reason to believe that this is not the case for SRRM4 and its worm ortholog RSR-2. The C terminus of SRRM4 appears to be essential for its microexon-stimulating role, and this domain is not present in worm

RSR-2^10^. We therefore speculate that stimulation of microexons in the *C. elegans* nervous system is accomplished by other sequence-specific RNA binding proteins that in turn recruit PRP-40. For example, the splicing factor RBFOX, which regulates some mammalian microexons^4^, or other neuron-specific splicing factors^20,21^, might facilitate neuronal microexon splicing.

One of the strongest splicing changes upon PRPF40A knockdown is loss of a poison exon in the *Luc7l* gene. We propose that this constitutes a compensatory homeostatic mechanism by which loss of PRPF40A leads to an increase in its spliceosomal binding partner LUC7L. Such compensatory mechanisms are often observed between paralagous RNA binding proteins, and indeed such a compensatory mechanism is observed between *Luc7l* and its paralog *Luc7l2*^14,15^. Here we extend this phenomenon to also apply to functionally related but evolutionarily unrelated factors. Another interesting link between PRPF40A and LUC7L is that they are both U1 snRNP-associated proteins, but have restricted effects on sub-classes of alternative splicing events. For PRPF40A, regulation is specific to microexons and small exons. For LUC7L, regulation is specific to certain classes of 5’ splice sites ^15,22^. This suggests a degree of specialization within individual snRNPs such that different components are responsible for regulating different aspects of mRNA splicing.

## MATERIALS AND METHODS

### N2a cell maintenance and harvesting

N2a (Neuro-2a) cells obtained from ATCC (CCL-131) were grown in OptiMEM medium plus DMEM, high glucose with L-glutamine, with 5% FBS and penicillin/streptomycin at 37°C and 5% CO2. Cells were grown in biological triplicates.

For gene-specific knockdown, cells were transfected with 10 nM pools of gene-specific siRNA (siGENOME, Dharmacon) using RNAiMax, according to manufacturer recommendations (Life Technologies). Cells were harvested 72 hours after transfection for RNA-Seq analysis. RNA was extracted using Tri reagent and ZYMO Direct-zol RNA miniprep kits.

### RNA-Seq library preparation and analysis

Libraries were prepared using NEBnext Ultra II polyA-enriched kits for Illumina sequencing, followed by quality control measurements via Qubit and Agilent Tapestation. 150 bp paired-end reads were generated on an Illumina HiSeq 2000 short-read sequencer, and reads were mapped to the mouse genome (GRCm38) with STAR^23^ (version 2.5.3a). Each condition was sequenced in biological triplicate, with an average of 41.5 million paired-end reads per replicate, and an average of 74.1% uniquely-mapped reads per replicate. All sequencing reads and data are deposited at NCBI Gene Expression Omnibus (GEO) database (Accession GSE267257). Previously-published *prp-40* mutant RNA Seq data from *C. elegans* is available at the NCBI SRA (Accession PRJNA684142).

Gene-specific counts were tabulated for each sample using HTSeq^24^ (version 0.9.1) and statistically-significant differentially expressed transcripts were identified with DESeq2^25^ (version 1.36.0).

Differential alternative splicing analysis was carried out using the Junction Usage Model ^26^ (JUM 2.0.2). For visualization of aligned reads, bam files were generated by samtools and subsequently sashimi plots were generated by the Integrative Genomics Viewer (IGV).

## REFERENCES

1. Ramírez-Sánchez, O., Pérez-Rodríguez, P., Delaye, L. & Tiessen, A. Plant Proteins Are Smaller Because They Are Encoded by Fewer Exons than Animal Proteins. Genomics Proteomics Bioinformatics 14, 357–370 (2016).

2. Berget, S. M. Exon recognition in vertebrate splicing. J. Biol. Chem. 270, 2411–2414 (1995).

3. Dominski, Z. & Kole, R. Selection of splice sites in pre-mRNAs with short internal exons. Mol. Cell. Biol. 11, 6075–6083 (1991).

4. Li, Y. I., Sanchez-Pulido, L., Haerty, W. & Ponting, C. P. RBFOX and PTBP1 proteins regulate the alternative splicing of micro-exons in human brain transcripts. Genome Res. 25, 1–13 (2015).

5. Irimia, M. et al. A highly conserved program of neuronal microexons is misregulated in autistic brains. Cell 159, 1511–1523 (2014).

6. Gonatopoulos-Pournatzis, T. & Blencowe, B. J. Microexons: at the nexus of nervous system development, behaviour and autism spectrum disorder. Curr. Opin. Genet. Dev. 65, 22–33 (2020).

7. Quesnel-Vallières, M., Irimia, M., Cordes, S. P. & Blencowe, B. J. Essential roles for the splicing regulator nSR100/SRRM4 during nervous system development. Genes Dev. 29, 746–759 (2015).

8. Calarco, J. A. et al. Regulation of vertebrate nervous system alternative splicing and development by an SR-related protein. Cell 138, 898–910 (2009).

9. Choudhary, B., Marx, O. & Norris, A. D. Spliceosomal component PRP-40 is a central regulator of microexon splicing. Cell Rep. 36, 109464 (2021).

10. Torres-Méndez, A. et al. A novel protein domain in an ancestral splicing factor drove the evolution of neural microexons. Nat. Ecol. Evol. 3, 691–701 (2019).

11. Li, X. et al. A unified mechanism for intron and exon definition and back-splicing. Nature 573, 375–380 (2019).

12. Ruan, J. et al. TreeFam: 2008 Update. Nucleic Acids Res. 36, D735–740 (2008).

13. Ester, C. & Uetz, P. The FF domains of yeast U1 snRNP protein Prp40 mediate interactions with Luc7 and Snu71. BMC Biochem. 9, 29 (2008).

14. Jourdain, A. A. et al. Loss of LUC7L2 and U1 snRNP subunits shifts energy metabolism from glycolysis to OXPHOS. Mol. Cell 81, 1905-1919.e12 (2021).

15. Kenny, C. J. et al. LUC7 proteins define two major classes of 5’ splice sites in animals and plants. Preprint at 10.1101/2022.12.07.519539 (2022).

16. Ni, J. Z. et al. Ultraconserved elements are associated with homeostatic control of splicing regulators by alternative splicing and nonsense-mediated decay. Genes Dev. 21, 708–718 (2007).

17. Lareau, L. F. & Brenner, S. E. Regulation of splicing factors by alternative splicing and NMD is conserved between kingdoms yet evolutionarily flexible. Mol. Biol. Evol. 32, 1072–1079 (2015).

18. Spellman, R., Llorian, M. & Smith, C. W. J. Crossregulation and functional redundancy between the splicing regulator PTB and its paralogs nPTB and ROD1. Mol. Cell 27, 420–434 (2007).

19. Gonatopoulos-Pournatzis, T. et al. Genome-wide CRISPR-Cas9 Interrogation of Splicing Networks Reveals a Mechanism for Recognition of Autism-Misregulated Neuronal Microexons. Mol. Cell 72, 510-524.e12 (2018).

20. Norris, A. D., Gracida, X. & Calarco, J. A. CRISPR-mediated genetic interaction profiling identifies RNA binding proteins controlling metazoan fitness. eLife 6, e28129 (2017).

21. Taylor, M., Marx, O. & Norris, A. TDP-1 and FUST-1 co-inhibit exon inclusion and control fertility together with transcriptional regulation. Nucleic Acids Res. 51, 9610–9628 (2023).

22. Puig, O., Bragado-Nilsson, E., Koski, T. & Séraphin, B. The U1 snRNP-associated factor Luc7p affects 5’ splice site selection in yeast and human. Nucleic Acids Res. 35, 5874–5885 (2007).

23. Dobin, A. et al. STAR: ultrafast universal RNA-seq aligner. Bioinforma. Oxf. Engl. 29, 15–21 (2013).

24. Anders, S., Pyl, P. T. & Huber, W. HTSeq--a Python framework to work with high-throughput sequencing data. Bioinforma. Oxf. Engl. 31, 166–169 (2015).

25. Love, M. I., Huber, W. & Anders, S. Moderated estimation of fold change and dispersion for RNA-seq data with DESeq2. Genome Biol. 15, 550 (2014).

26. Wang, Q. & Rio, D. C. JUM is a computational method for comprehensive annotation-free analysis of alternative pre-mRNA splicing patterns. Proc. Natl. Acad. Sci. U. S. A. 115, E8181–E8190 (2018).

